# BioCAT: a novel tool to search biosynthetic gene clusters producing nonribosomal peptides with a known structure

**DOI:** 10.1101/2021.09.13.460047

**Authors:** D.N. Konanov, D.V. Krivonos, E.N. Ilina, V.V. Babenko

## Abstract

**Motivation:** Nonribosomal peptides are a class of secondary metabolites synthesized by multimodular enzymes named nonribosomal peptide synthetases and mainly produced by bacteria and fungi. It has been shown that non-ribosomal peptides have a huge structural and functional diversity including antimicrobial activity, therefore, they are of increasing interest for modern biotechnology. Methods such as NMR and LC-MS/MS allow to determine a peptide structure precisely, but it is often not a trivial task to find natural producers of them. Today, the search is usually performed manually, mostly with tools such as antiSMASH or Prism. However, there are cases when potential producers should be found among hundreds of strains, for instance, when analyzing metagenomes data. Thus, the development of automated approaches is a high-priority task for further nonribosomal peptides research.

**Results:** We developed BioCAT, a two-side approach to find biosynthetic gene clusters which may produce a given nonribosomal peptide when the structure of interesting nonribosomal peptide has already been found. Formally, BioCAT unites the antiSMASH software and the rBAN retrosynthesis tool but some improvements were added to both gene cluster and peptide chemical structure analyses. The main feature of the method is an implementation of position specific score matrix to store specificities of nonribosomal peptide synthetase modules, which has increased the alignment quality in comparison with more strict approaches developed earlier. An ensemble model was implemented to calculate the final alignment score. We tested the method on a manually curated nonribosomal peptides producers database and compared it with a competing tool called GARLIC. Finally, we showed the method applicability on several external examples.

**Availability:** BioCAT is available on the GitHub repository or via pip

## 1 Introduction

Nonribosomal peptides (NRPs) are secondary metabolites produced by a wide range of taxa, such as bacteria, fungi, plants and even animals [1].. Biosynthesis of NRPs in cells is provided by multidomain enzymes named nonribosomal peptide synthetases (NRPS) in an iterative way (Figure 1). Each functional module of NRPS generally consists of three domains: the adenylation domain (A-domain) providing the activation of the substrate using ATP, the peptidyl-carrier domain (PCP-domain) binding the substrate to the 4’-phospho-pantethine group of PCP-domain, and the condensation domain (C-domain) catalyzing the amide bond or, in a number of cases, the ester bond [2] formation between substrates from the current and previous modules (Fig. 1). Additionally, the last module of NRPS should contain the thioesterase domain (TE-domain) which hydrolyzes the thioester bond realizing the biosynthesis product. It was shown that the modules substrate specificity mostly provided by A-domains [3] but some reports describing C-domain specificity also were published [4, 5].

**Figure 1:**
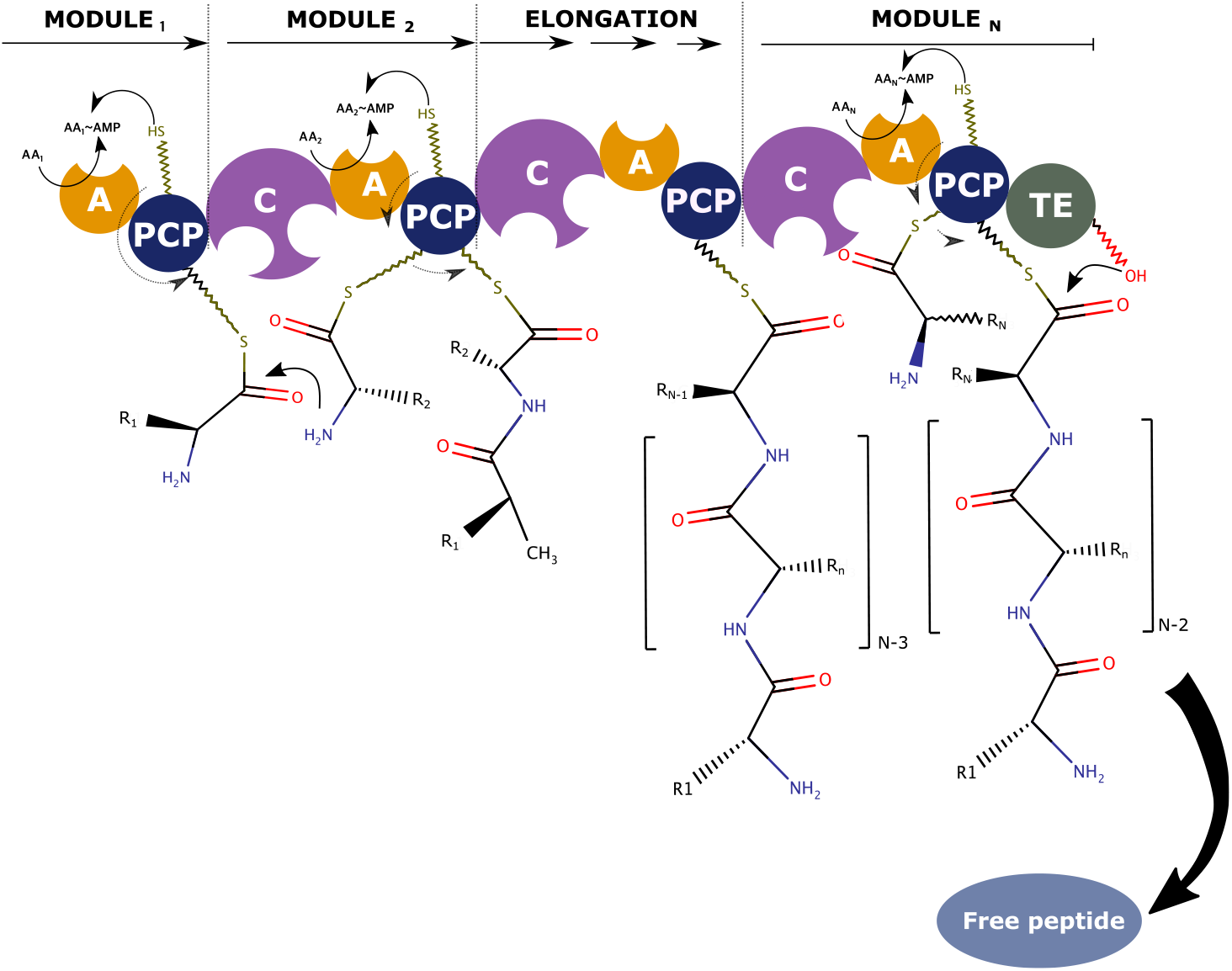
Typical NRP biosynthesis scheme. The adenylation domain (A-domain) from Module 1 activates the first aminoacid using ATP. Next, the neighbouring peptidyl-carrier protein domain (PCP-domain) forms the thioester bond with the activated aminoacid. Simultaneously, the same process occurs in the Module 2. After aminoacids have been bound to corresponding PCPs, the condensation domain (C-domain) from Module 2 catalyzes peptide bond formation between them. The process is iteratively repeated until a thioesterase domain (TE-domain) from the last module realizes the biosynthesis product.

The unique biosynthesis scheme has led to the tremendous diversity in the molecular structure of NRPs in comparison with ribosome-synthesized peptides, firstly, due to the possibility to combine both proteinogenic and non-proteinogenic substrates as well as to modify monomers by hydroxylation, halogenation, epimerization and other ways simultaneously with the biosynthesis process. Moreover, joint work of different enzymes allows to build more complex structures such as NRP-polyketide hybrids [6], NRPs containing *β*-lactam ring [7], cyclic depsipeptides [8] and others.

An accurate prediction of potential producers of a given NRP is not a trivial task even when the NRP structure is known because of two main reasons. On the one hand, A-domains specificity prediction is based on a slightly small NRP producers dataset available these days. Thus, existing tools such as SVM-based NRPSPredictor 2 [9] and an ensemble method called SANDPUMA [10] have been trained on less than one hundred manually annotated A-domains which seems insufficient for accurate specificity prediction, mainly because of the high monomers variety. The second problem is related to the complexity of accurate NRP retrosynthesis. In addition, a number of NRP structures is synthesized in non-iterative schemes including dimerization of peptide fragments (Type B biosynthesis pathway [11]) or use of one NRPS module more than one time during the biosynthsis (Type C biosynthesis pathway [11]).

Here, we present BioCAT (Biosynthesis Cluster Analysis Tool), a new tool which allows to find producers of a given NRP, using as the input a SMILES-formatted chemical structure and the genome of the potential producer in the FASTA format. Formally, the method unites antiSMASH 6 [12] biosynthetic gene cluster (BGC) predictions and the rBAN [13] retrosynthesis tool, but there are a number of improvements added to both gene cluster and chemical structure analyses. Firstly, we developed a PSSM-based approach to align NRP and BGC. Secondly, we implemented the retrosynthesis model which generates not just monomers but probable pathways of synthesis which we named core peptide chains. It should be noted, that the tool is designed to analyse only prokaryotic genomes because of insufficient size of fungal NRP producers data.

To validate our model, we checked the quality of the full pipeline on the manually curated dataset of all known NRP/producer pairs using shuffle-split cross-validation. In addition, we showed the applicability of BioCAT on several external data, including complete genomes as well as draft ones. Finally, we compared the BioCAT pipeline with the GARLIC tool [14] which has a similar functionality.

## 2 Materials and methods

### 2.1 Database collection

BGC annotations and corresponding chemical structures for 426 known NRPs were collected from MIBiG [15] database. To ensure consistency of annotations all BGCs were re-annotated using antiSMASH 6 [12]. 1675 A-domain sequences with known specificity were extracted (full list of used sequences is available in Supplementary Data, A-domains table). To check genome-level applicability 164 prokaryotic genomes containing known BGCs were collected from NCBI (full list of collected NRP-genome pairs is available in Supplementary Data, AllProducers table).

### 2.2 NRP retrosynthesis

In BioCAT, the NRP structure processing consists of two main parts: the monomers identification by rBAN [13] and the extracting of peptide fragments which we named core peptide chains. Additionally, there are a number of features improving the parsing of NRP chemical structures such as cycles solving and searching of inner fragments which are probably synthesized in non-classic ways (e.g. Type B or Type C biosynthesis pathways).

#### 2.2.1 Core peptide chain(s) prediction

Firstly, the SMILES-formatted NRP structure is processed by the rBAN software [13] (discovery mode is enabled). Next, each bond in the resulting monomeric graph (Fig. 2, 1) is checked to be the peptide bond. If some bond is not peptide, it will be removed from the graph. Thereafter, we have a number of distinct peptide fragments which are supposed to be synthesized in a linear way during the biosynthesis process. Next, each monomer is checked to be an *α*-aminoacid strictly, and all non-aminoacid monomers are removed from the fragments. If some fragments remain to be cyclic, the algorithm will hydrolyze these fragments in all possible ways to get all possible linear monomeric sequences. Simultaneously, the algorithm checks that the considered product can be synthesized in the Type B or Type C ways (if this option is enabled) and generates additional monomeric sequences modified according to these biosynthesis types. Thus, on this stage, we have a number of linear peptide fragments consisting only of *α*-aminoacids bounded only by peptide bonds (Fig. 2, 2). To generate the product sequences which will be aligned against a given PSSM, these fragments are concatenated in all possible ways. If the length of concatenates is less than the size of the PSSM, a required number of gaps will be added between concatenated fragments. Gaps in the concatenates are assigned as *nan* (Fig. 2, 3A and 3B). In the further sections, we will call these concatenates core peptide chains (CPCs).

**Figure 2:**
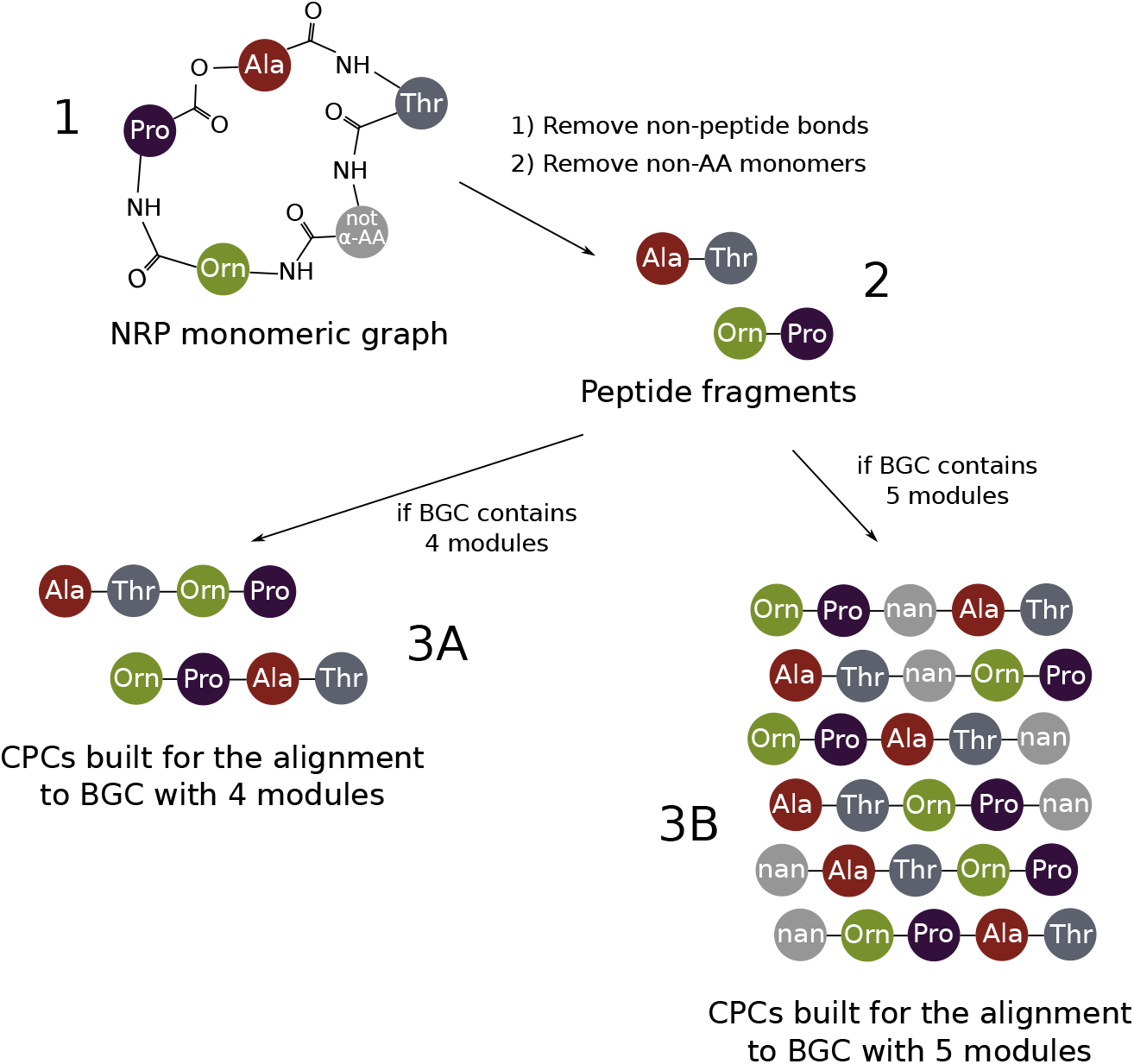
Combinatorial approach to align NRP structure against BGC. Firstly, the monomeric graph generated by rBAN (1) is cut along all bonds which were recognized as non-peptide. Simultaneously, all monomers which were not recognized as *α*-aminoacids are removed from the graph. Resulting peptide fragments (2) can be combined in different ways which depend on the size of BGC with which the current alignment is performed. If the number of modules in the BGC is the same as the sum number of monomers in the peptide fragments, these fragments will be just rearranged in all possible ways to generate core peptide chains (CPCs) (3A). If the number of modules in the BGC is more than the sum number of monomers in the peptide fragments, gaps assigned as *nan* will be added to core peptide chains to all possible positions (3B).

**Figure 3:**
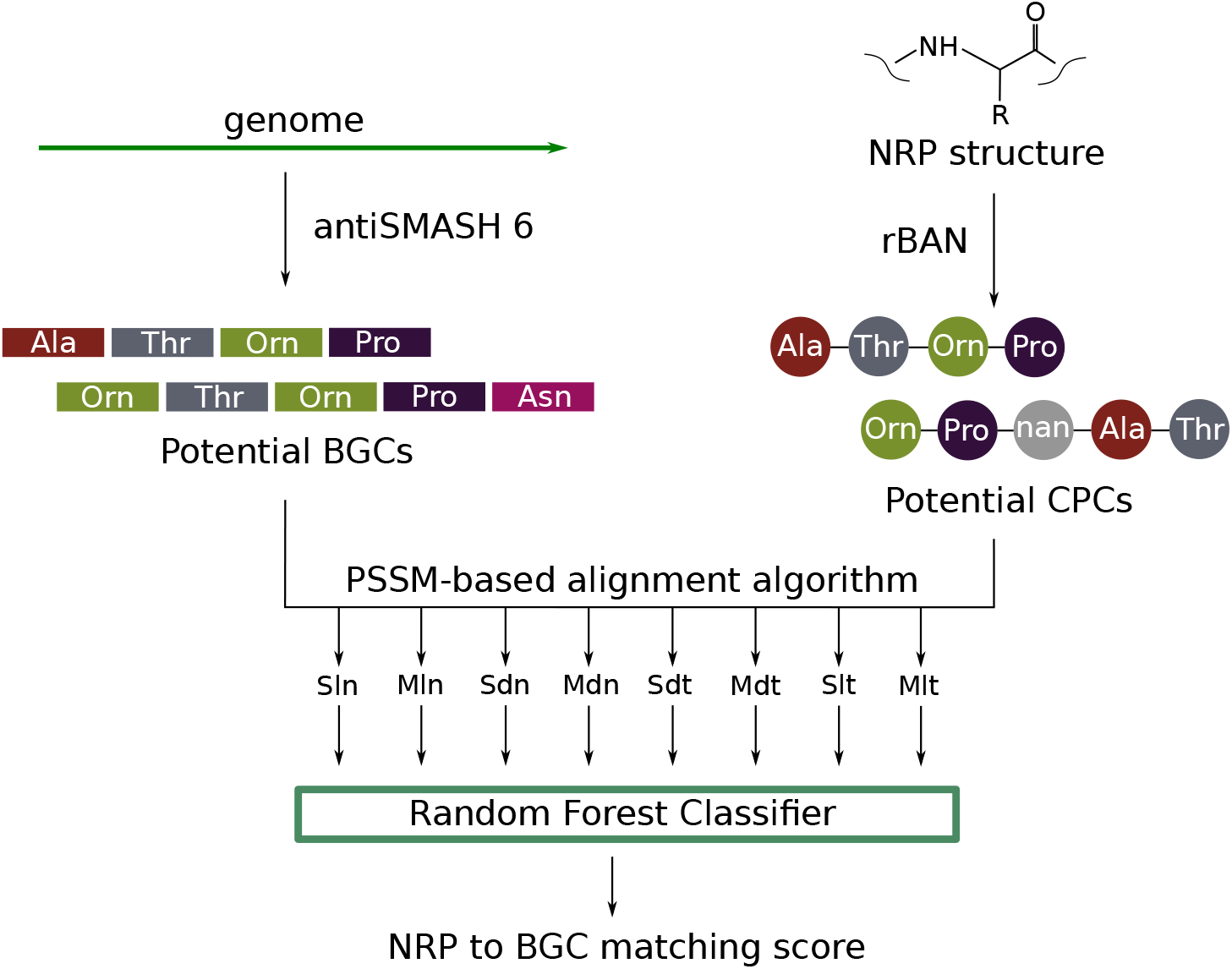
Principal scheme of the BioCAT pipeline. First, the input genome is processed by antiSMASH 6 and the NRP structure is processed by rBAN. Next, all potential biosynthesis gene clusters (BGCs) are aligned against all possible core peptide chains (CPCs) built from the monomeric graph generated by rBAN. Alignment is performed using eight different variants of the alignment score definition (Sln, Mln, Sdn, etc). Finally, for each successful matching, these scores are processed by the Random Forest Classifier, which generates the final matching score distributed from 0 to 1 and the binary matching score.

### 2.3 BGC analysis

#### 2.3.1 Profile Hidden Markov Models construction

The most common substrates were chosen such that for each of them there were at least 10 A-domain sequences in the database. Only these sequences were used in the further analysis. Profile HMMs construction were carried out as follows. Suppose that we have a number of A-domain sequences for *i*-th substrate. Firstly, we use these sequences as the base to build a profile HMM for *i*-th substrate using HMMER3 [16]. Next, we take all A-domains sequences which are known not to have the specificity to *i*-th substrate and align them against this profile HMM for *i*-th substrate. Therefore, we have a number of alignment scores which we have named the negative background for *i*-th substrate (*NB_i_*). In this pipeline, E-values given by HMMER3 were used as the alignment scores.

#### 2.3.2 PSSM construction

Suppose that *X* = (*x*_1_,*x*_2_,…,*x*_N_) is an NRPS modules sequence with unknown specificity and there are *N* distinct A-domains already annotated. Consider *i*-th A-domain sequence *X_i_* and profile HMM for *j*-th substrate. Raw specificity score for this pair is *E*-value of the sequence to HMM alignment. Let’s assume that it equals *starget*. We defined the relative specificity score *g_ji_* as:

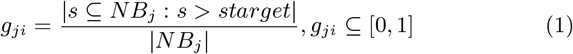

where *NB_i_* is the negative background for *j*-th substrate. In simple words, the closer this number is to one, the greater the chance that *i*-th A-domain has a specificity to *j*-th substrate.

After this procedure is carried out for all *N* A-domains and *S* substrates, we get the following matrix:

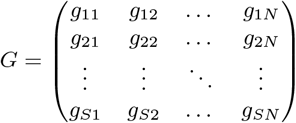

where *S* is the number of possible substrates, *N* is the number of modules in the BGC, *g_ji_* is the chance that *i*-th A-domain has a specificity to *j*-th substrate.

In bioinformatics, *G* is a classic example of position-specific score matrix (PSSM) which can be used as an alignment template.

### 2.4 Alignment process

As the input for the alignment we should take one core peptide chain vector *P* = (*p*_1_, *p*_2_,…, *p_N_*) and one PSSM matrix *G* with dimension (*S, N*).

#### 2.4.1 Alignment score definition

An ensemble model has been developed for an efficient NRP to BGC alignment. First, we implemented two ways to compute raw alignment scores:

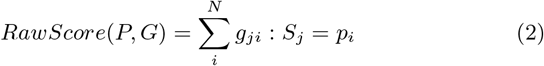

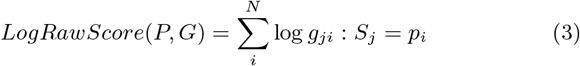

In both cases, if *p_i_* equals *nan*, zero score will be added for *i*-th module.

The linear sum (Eq. 2) was the most intuitive and had a satisfactory prediction quality (F1-score = 0.596) but returned a lot of false positive matches. The logarithmic sum (Eq. 3) turned out to be more specific due to the higher influence of too small values in the PSSM on the final score. However, it had less general quality (F1-score = 0.574) compared with the linear approach.

Secondly, we build the monomers sequence *MaxSeq* = [*ms*_1_, *ms*_2_,…, *ms_N_*] by the following way:

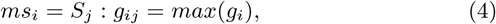

where *g_ij_* are elements of the aligned PSSM. There also were two options, how *MaXSeq* is built. The first is insertion of *nan* to the positions which contain *nan* in the core peptide chain sequence (replaced *MaxSeq*). Such operation significantly increases the method sensitivity but, again, leads to the increase in the false positive rate. If *nan* are not inserted to the *MaxSeq* (native *MaxSeq*) an absolute value of the raw alignment score tends to be much lower.

The absolute value of *RawScore* or *LogRawScore* depends on the core peptide chain length, the count of *nan* in the sequence and the nature of monomers included in the core peptide chain. To estimate the quality of an alignment, *I* randomly shuffled PSSM matrices are generated. The shuffling is performed in two different ways: by rows (intermodular shuffling) or by columns (intersubstrate shuffling). *I* was chosen to be 100 by default.

Next, after two *MaxSeq*-s and two types of shuffled matrices were formed, both native and replaced *MaxSeq* is aligned against each shuffled PSSM using both linear and logarithmic raw score calculation ways. Combining all possible computing options, we have eight arrays *F_k_* (*k* = 1, 2,…, 8) each of which contains *I* shuffled raw scores. Suppose that the observed core peptide chain was aligned to the non-shuffled PSSM with raw score equals *target*. We defined the relative alignment score for *k*-th method as:

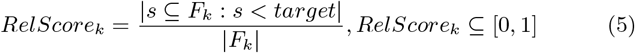

In other words, the relative alignment score shows the fraction of shuffled scores which are less than the non-shuffled score. Distributions of relative score for all individual models obtained on the positive dataset and negative control are shown on Supplementary Figure 1.

Finally, these eight relative scores are processed by the Random Forest model which generates the final score also distributed between 0 to 1. Values close to 1 can be considered as successful matches.

#### 2.4.2 Best match logic

As it was mentioned in the previous sections, a core peptide chain cannot be unambiguously determined in most cases. Due to this, the combinatorial approach was implemented to generate all possible peptide chains which can be matched to a current PSSM. The Random Forest model score is computed for each peptide chain variant independently and the highest score is chosen as the alignment result.

### 2.5 Output explanation

Despite only the highest alignment score influences the final alignment report, all combinatorial chain alignments are saved into the resulting file. Generally, the resulting file is a table consisting of the following columns:

- Chromosome name (1 column)
- Coordinates of BGC (2 column)
- Strand (3 column)
- Substance name (4 column)
- Cluster ID (5 column)
- Core peptide chain (6 column)
- Supposed biosynthesis type (e.g. Type A, B or C) (7 column)
- Sln, Mln, Sdn, Mdn, Sdt, Mdt, Slt, Mlt scores (8-15 columns)
- Probability of successful match for current alignment (16 column)
- Random Forest binary prediction (17 column)

8-15 columns of the resulting file contain scores returned by eight different individual models. Names of individual models describe their parameters as following:

- First letter means PSSM shuffling type (‘S’ is intersubstrate, ‘M’ is intermodular)
- Second letter means raw score calculation type (‘l’ is logarithmic, ‘d’ is linear)
- Third letter means *MaxSeq* processing option (‘n’ is with *nan* insertion, ‘t’ is without insertion)

### 2.6 Method validation

First, we generated 820 incorrect genome/NRP pairs to estimate the false positive rate of considered methods. The number of incorrect pairs was chosen to be 5 times more than the number of correct pairs to check the specificity and selectivity of the method more accurately. To validate the Random Forest classification model implemented in BioCAT, the database of 984 genome/NRP pairs (164 correct + 820 incorrect) was divided in 80:20 ratio on train and test sample respectively. The accuracy of matching was estimated using precision, recall, F1-score and MCC metrics. Additionally, a receiver operating curve (ROC) and a precision-recall (PR) curve were built using scores returned by the model. To estimate the stability of the model, the train/test splitting was performed randomly in 1000 iterations. Parameters used for the Random Forest model construction are available in the Supplementary Data, RFParameters table. OOB error curves and feature wights are shown on Supplementary Figures 2 and 3.

The method was compared with the GARLIC pipeline [14] which has a similar functionality. The latest versions of GRAPE (1.0.2) and PRISM (2.1.5) tools for which command-line versions were available were used. 164 genome sequences containing BGC with a known product were analysed by PRISM to locate BGCs. Retrosynthesis of chemical structures was performed by GRAPE. The same list of 984 correct and incorrect genome/NRP pairs was processed by the GARLIC. Next, relative scores returned by GARLIC also were divided on train and test samples in 80:20 ratio and used as the base for the linear classifier. Procedure was repeated 1000 times. The same classification accuracy metrics as for the BioCAT were calculated.

In the analysis, if any method did not return any possible alignments for a pair, the alignment score was assumed to be zero.

### 2.7 Used software and tools

MUSCLE 3.8.1551 [17] was used for multiple alignment. HMMER3 3.1b2 [16] was used to build profile HMMs. rBAN 1.0 [13] was used for the NRPs retrosynthesis. To locate biosynthesis gene clusters antiSMASH 6.0.0 [12] was used, with Prodigal 2.6.3 [18] as a gene finding tool. The random forest model was implemented using the Scikit-learn python library (v0.24.2) [19].

## 3 Results

### 3.1 Software description and availability

We have developed a tool that estimates the likelihood that a given non-ribosomal peptide synthetase, or more generally a given organism, is capable of producing a given NRP. The tool is available as a CLI program named BioCAT (Biosynthesis Cluster Analysis Tool) on GitHub (https://github.com/DanilKrivonos/BioCAT) or can be installed via pip. The required input files necessary for the analysis are a FASTA-formatted genome and SMILES-formatted NRP-structure. In the BioCAT pipeline, the genome sequence is analysed by antiSMASH 6 and the structure is characterized by rBAN, so, these programs are required to be installed. Additionally, it is possible to use pre-calculated antiSMASH or rBAN results in JSON-format.

Biosynthesis of NRPs can be carried out not only in the strict iterative way shown on Fig 1. We will use the NRP biosynthesis type notation proposed in [11], where the most common canonical iterative pathway is called Type A, and two additional variants of the NRP building called Type B and Type C are defined. The Type B pathway includes a formation of two or more identical NRP fragments catalyzed by the same NRPS or the same part of NRPS which will be bound in further biosynthesis stages. NRPs such as actinomycin D are shown to be synthesized in the Type B pathway [20]. The Type C biosynthesis variant shown for such NRPs as lugdunin [21] includes a consequent binding of two or more identical monomers to the growing peptide chain catalyzed by one NRPS module. In BioCAT, we implemented the support of both non-linear biosynthesis types.

We have found that the best producers prediction quality can be reached using ensemble approaches. We have implemented the random forest classifier model which computes the final alignment score using eight pre-scores generated by slightly different algorithms, which are described in details in the Materials and Methods section.

The result of BioCAT analysis is information about all possible NRP to BGC alignments generated in a combinatorial way. For each alignment the final alignment score is computed independently. Final scores returned by BioCAT are distributed from 0 to 1, where values close to one show that the given BGC is likely to code the NRP synthetase providing the biosynthesis of the given NRP and vice versa.

### 3.2 Method quality

A manually curated database consisting of full genomes of 164 known NRP chemical structures and corresponding NRP producers was collected. These genome/NRP pairs were analysed using the BioCAT and the GARLIC [14] algorithms, because, in our knowledge, GARLIC is the only tool, which has the same functionality and can be compared directly with the BioCAT pipeline. It should be noted that the GARLIC tool is based on another BGC prediction and NRP retrosynthesis software (PRISM and GRAPE respectively), so, the differences in the methods quality can be caused not only by the differences between considered alignment algorithms but also by the accuracy of these stages. To estimate the false discovery rate, randomly generated genome/NRP pairs were aligned by both the BioCAT and GARLIC algorithms. The number of generated incorrect genome/NRP pairs was chosen to be 5 times more than the number of correct pairs. Total, 984 genome/NRP pairs (164 correct + 820 incorrect) were analysed. The list of pairs was the same for both used methods. In the analysis, if any method did not return any possible alignments for a pair, the alignment score was assumed to be zero.

We have estimated the accuracy of the methods using recall, precision, F1-score and MCC metrics (Table 1). After 1000 iterations of random 80:20 train/test splitting, BioCAT had a higher mean recall (0.710 and 0.363 for BioCAT and GARLIC respectively), but GARLIC was shown to be more precision (0.556 and 0.766 respectively). Integral classification metrics such as F1-score and MCC also were higher on the BioCAT results. Additionally, a ROC curve and a precision-recall (PR) curve were built on the raw scores returned by both methods (Fig. 4). BioCAT showed a higher PR AUC value (GARLIC PR AUC = 0.62, BioCAT ROC AUC = 0.68). Average ROC AUC values were close (0.89 in both cases).

**Figure 4:**
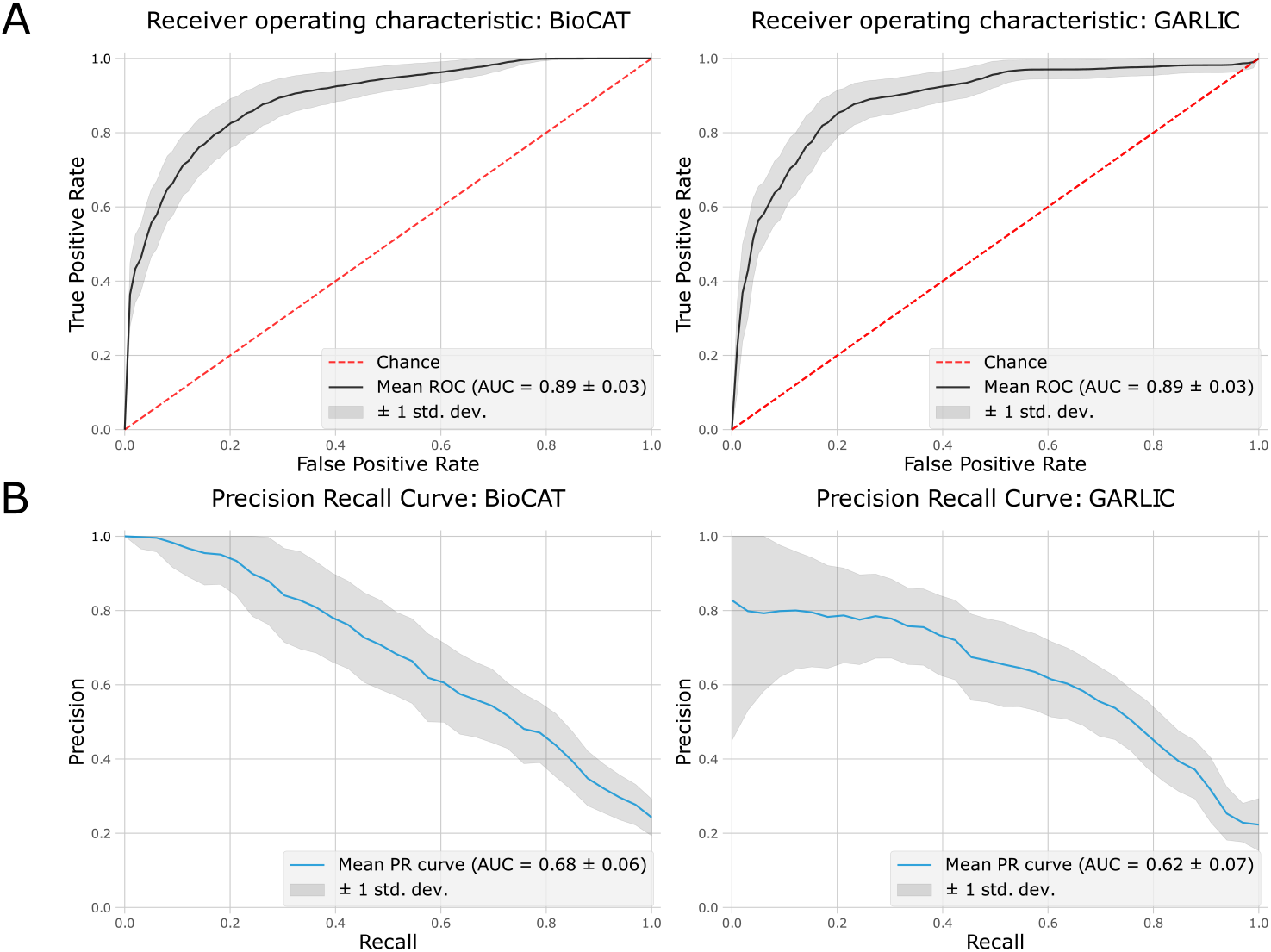
Performance of BioCAT compared with the GARLIC tool. Gray area shows the standard deviation of the y axis values during 1000 independent train/test splittings. A) Receiver operating characteristic curves obtained on test samples using BioCAT and GARLIC methods respectively. B) Recallprecision curves obtained on test samples using BioCAT and GARLIC methods.

**Table 1:** BioCAT performance compared with the GARLIC tool.

| method | recall | precision | F1-score | MCC | Mean time consumption, s |
| --- | --- | --- | --- | --- | --- |
| BioCAT | 0.710 | 0.556 | 0.619 | 0.540 | 332 |
| GARLIC | 0.363 | 0.766 | 0.487 | 0.468 | 527 |

Next, we trained the Random Forest model on all data and performed new analysis on the full dataset. The resulting ROC AUC was 0.93 which was close to the value obtained on test datasets and might indicate that the model had not been heavily overfitted.

### 3.3 Method benchmarking

During the analysis, time consumption of both considered methods was measured. We found that in both methods used the BGC detection was the limiting stage. The total time taken by the BioCAT pipeline was 330 seconds per NRP to genome alignment, which was faster than the full GARLIC pipeline, which averaged 527 seconds to run.

Additionally, we have tested how the alignment score depends on the number of shuffling iterations. Satisfactory convergence of the results was achieved at 100 iterations (Pearson’s correlation coefficient = 0.92), so this value was chosen by default (Supplementary Figure 4).

### 3.4 Method application

#### 3.4.1 Search for potential producers of a given NRP

After the model was trained and validated, a few external genome/NRP pairs were processed to show the applicability of the method. Laterocidin [22], thanamycin [23] and mutanobactin [24] which were not included into the curated dataset were aligned against their producers and a number of related genomes from the same genera. Laterocidin (Fig. 5A) was successfully aligned against its producer *Brevibacillus laterosporus* LMG 15441 with a relative score of 0.81. At the same time all alignments against other *Brevibacillus* strains returned relative alignment scores less than 0.5 (Supplementary data, Laterocidine_test). Interestingly, thanamycin (Fig. 5B) had the successful matching score not only with its own producer *Pseudomonas fluorescens* DSM 11579, but with four other strains of *Pseudomonas* (Supplementary data, Thanamycin_test). One of them, *Pseudomonas sp*. 11K1, has been shown to produce brasmicin, an NRP homologous to thanamicin [25]. Thus, others can also be considered as potential producers of NRPs with a similar monomeric sequence.

**Figure 5:**
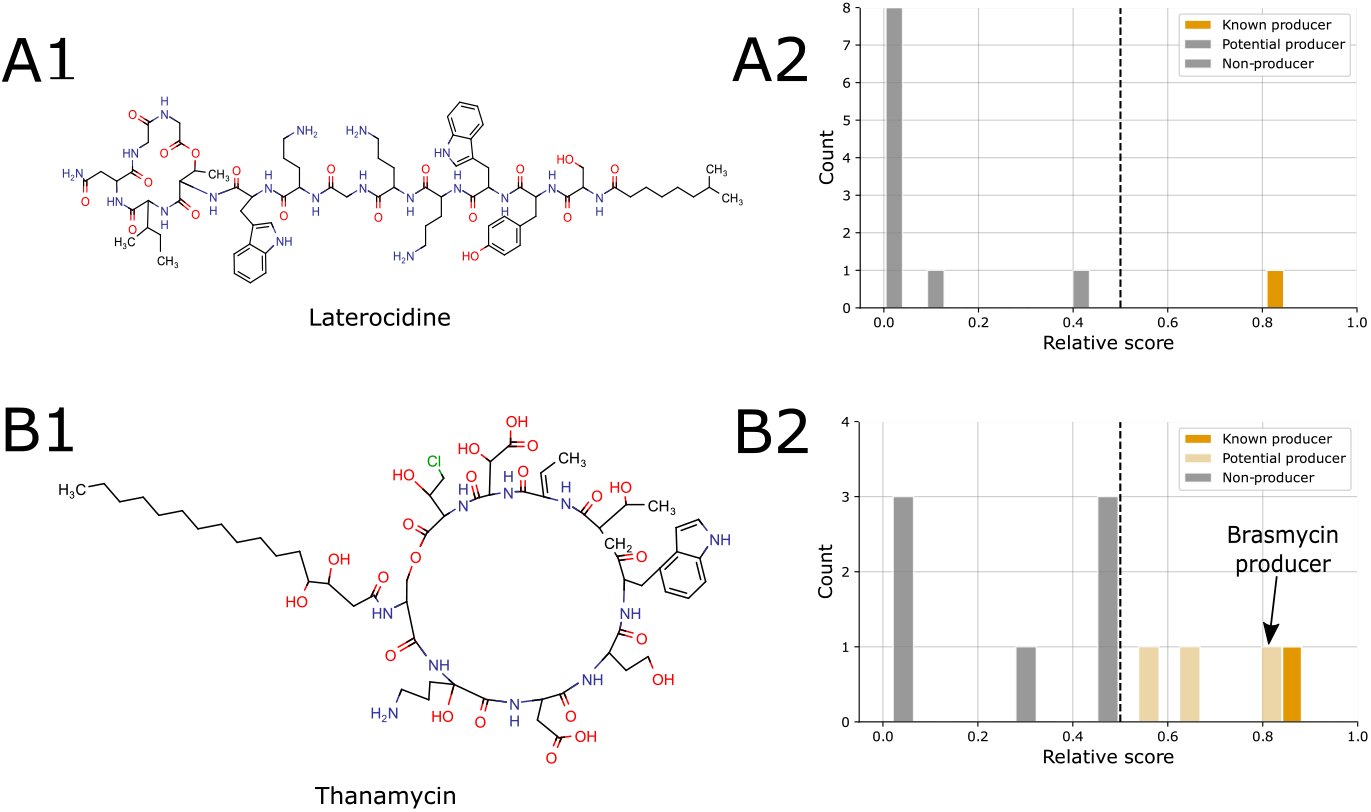
Applicability of BioCAT for the identification of potential producers of a given NRP. A) Laterocidine chemical structure (A1) was aligned against its natural producer *Brevibacillus laterosporus* LMG 15441 and 10 close *Brevibacillus* strains. Only the producer (A2, orange bar) had the alignment score higher than 0.5. B) Thanamycin (B1) was aligned against its producer *Pseudomonas fluorencens* DSM 11579 (B2, orange bar) and 10 other *Pseudomonas* strains. The producer was successfully aligned with the resulting score of 0.86. In addition, there were three *Pseudomonas* strains which were assigned as potential producers of thanamycin (B2, light orange bars). *Pseudomonas sp*. 11K1 strain which had the highest alignment score was described earlier as a producer of brasmycin, NRP homologous to thanamycin.

Mutanobactin is one of the NRPs produced by *Streptococcus mutans*. In a recent work [26], the authors described in detail biosynthetic gene clusters in 17 different strains of *Streptococcus mutans* isolated from dental plaque. BGC responsible for the biosynthesis of mutanobactin was found in three strains (SA41, T4, 21). Using BioCAT, we found the same three strains as potential producers of mutanobactin with relative scores higher than 0.97. At the same time, all other strains did not have any successful matches. Additionally, we collected 21 different *Streptococcus mutans* complete genomes available in the RefSeq database and aligned them against the mutanobactin structure. *Streptococcus mutans* UA159 strain which had been described earlier as a producer of mutanobactin [24] had a relative score 0.97. Moreover, 7 additional strains (Supplementary data, Mutanobactin_test) were shown to have similar biosynthesis clusters.

## 4 Discussion

We have developed a new PSSM-based approach for NRP structure to biosynthesis gene cluster alignment and implemented it as a CLI tool called BioCAT (Biosynthesis Cluster Analysis Tool). In general, this tool is designed for search of potential producers of a given non-ribosomal peptide among a number of genomes, but also can be applied for solving the reversed task when a user is interested in searching of the most likely products which can be synthesized by a given organism. In the BioCAT pipeline, antiSMASH 6 [12] and rBAN [13] functionalities were united. In our knowledge, these tools are the most accurate in the context of BGC annotation and NRP retrosynthesis respectively, so, due to this they were chosen to be implemented in the pipeline.

During the BioCAT development, we were trying to avoid complex machine learning methods to store the transparency of the results interpretation. However, we found that different ways to define the alignment score provide either a high specificity or high selectivity but not both. Observing individual models results, we supposed that an ensemble model is capable of providing a higher accuracy than individual ones. We implemented a Random Forest Classifier combining eight slightly different methods to calculate the relative alignment score and showed its efficiency using common classification quality metrics. Additionally, we compared our method with the GARLIC pipeline [14] which, in our knowledge, is the only tool which has a similar functionality. NRP to BGC matching quality obtained on the BioCAT results turned out to be higher than the GARLIC quality.

The method we developed has a number of limitations, mainly related to the quality of NRP chemical structures retrosynthesis. The rBAN tool is able to determine a wide range of substrates but some unusual chemical modifications such as acylation of proline lead to the appearance of excessive unrecognized elements in a molecular graph. Moreover, some chemical features such as fatty acid residues or poly-ketide fragments are not used in the PSSM construction and are not taken into account during the processing. Also, in NRP biosynthesis, some condensation domains are known to be able to form not peptide bonds but ester ones [2]. In these cases, peptide chains will be restricted at the ester bond and resulting fragments will be combined more aggressively which can increase the chance of false positive results. Generally, if peptide chains generated by the model include too many substrates assigned as *nan*, we recommend users to try to simplify the chemical structure of the interesting NRP manually before the analysis, e.g. to remove modifications from the substrates.

The BGC prediction stage also has some drawbacks. The main is the lack of formal rules to define edges of biosynthetic gene clusters. For example, some gene clusters such as nunamicine/nunapeptine BGC from *Pseudomonas sp*. Ln5 encode two different NRPSs located close to each other because of regulatory reasons [27]. Fortunately, these NRPSs are encoded in different DNA strands, so, in BioCAT we implemented additional fragmentation of clusters based on the strand direction. However, there are cases, for example, himastatine biosynthesis cluster from *Streptomyces himastatinicus* ATCC 53653, when genes located in both DNA strands are responsible for a biosynthesis of only one NRP product [28].

Despite drawbacks described above, BioCAT showed a satisfactory matching accuracy and can be useful for a high-throughput exploratory analysis of genomic data to identify possible producers of an NRP of interest or structures homologous to it. Going forward, the method can be improved in several ways. First, we are planning to include to the model information about additional gene cluster domains such as halogenation and hydroxylation which may increase the specificity of the alignment algorithm. Secondly, core peptide chains generated during the BioCAT analysis often contain non-recognized substrates assigned as *nan*, so, these unrecognized peptide chain positions can be represented in the same PSSM way, using special metrics such as Tanimoto chemical similarity score. However, it can significantly increase the model complexity, so, we decided not to implement it in this version due to insufficient size of the NRP library available nowadays.

Simplification and unification of genomic data processing is becoming more important with the intensive development of sequencing technologies. Thus, the more massive genomes are sequenced, the more time is consumed to perform an accurate prediction of NRP producers among them manually using antiSMASH or Prism software. The authors do not declare that the BioCAT tool can completely replace manual BGC annotation but hope that it will help to automatize the preliminary genomic data observation to narrow down a list of possible producers of a given NRP.

## 5 Conclusion

We have developed a novel tool, called BioCAT, which has united the antiSMASH and the rBAN pipelines and allows to find potential producers of a given NRP. During the work, BioCAT was shown to be slightly faster and more precise in comparison with the GARLIC tool published earlier. The applicability of the method was additionally shown on several external data.

## Acknowledgements

The authors thank the Center for Precision Genome Editing and Genetic Technologies for Biomedicine, Federal Research and Clinical Center of Physical-Chemical Medicine of Federal Medical Biological Agency for providing computational resources for this project.

