## Supplementary material for "BioCAT: a novel tool to search biosynthetic gene clusters producing nonribosomal peptides with a known structure": Supplemetary Figures

### BioCAT: PSSM-based algorithm to search biosynthetic gene clusters producing nonribosomal peptides with known structure.
### **Supplemetary Materials.**

Konanov D.N. Krivonos D.V. Babenko V.I.
Ilina E.N.

September 13, 2021

#### 1 Supplementary Tables

| method | recall | precision | F1-score | MCC | Mean time consupmtion, s |
|---|---|---|---|---|---|
| BioCAT | 0.710 | 0.556 | 0.619 | 0.540 | 332 |
| GARLIC | 0.766 | 0.363 | 0.487 | 0.468 | 527 |

Supplementary Table 1: BioCAT performance compared with the GARLIC tool.

#### 2 Supplementary Figures

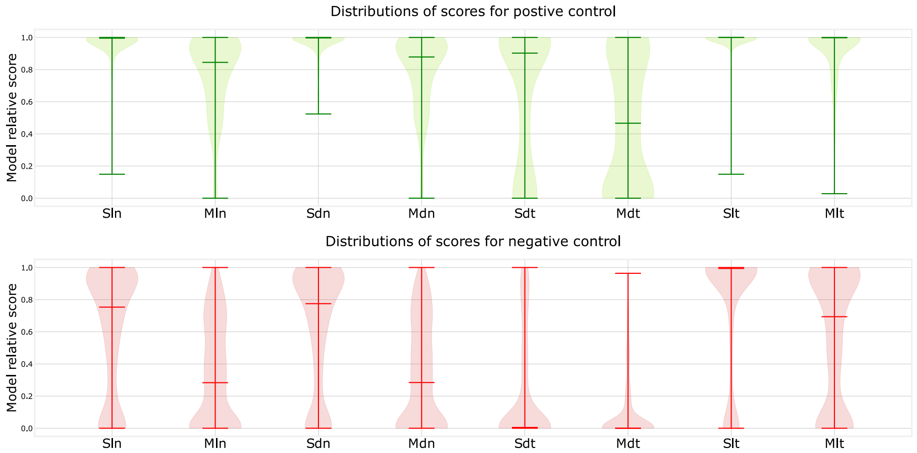


Supplementary Figure 1: Distributions of relative scores for each individual alignment model. Green violins show the positive NRP/genome pairs, red ones show negative control dataset.

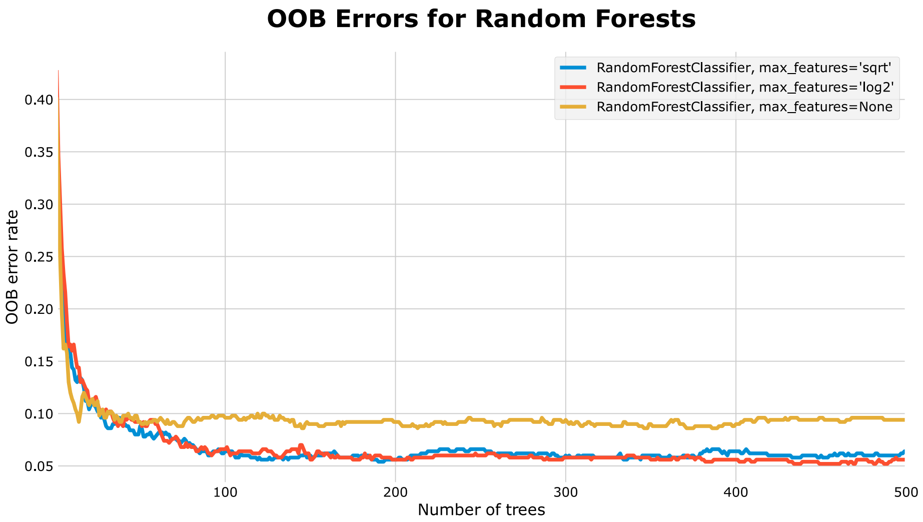


Supplementary Figure 2: OOB error curves obtained during Random Forest parameters optimization

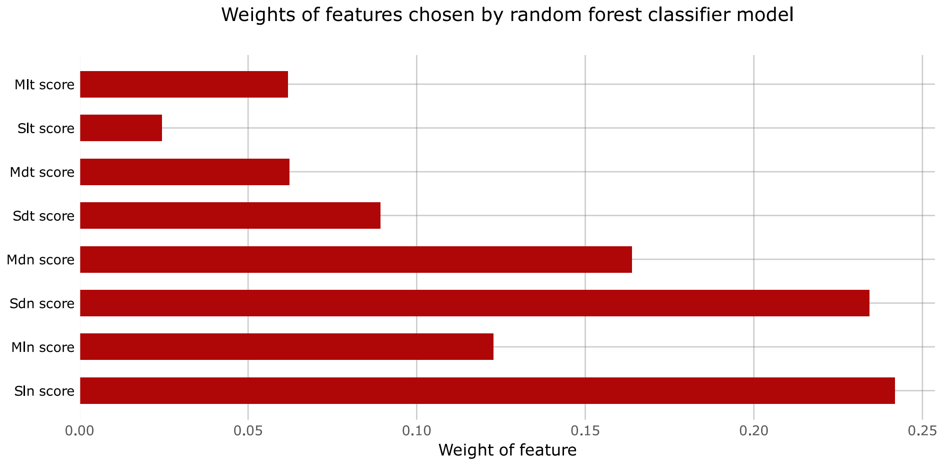


Supplementary Figure 3: Features weights in the Random Forest model implemented in BioCAT

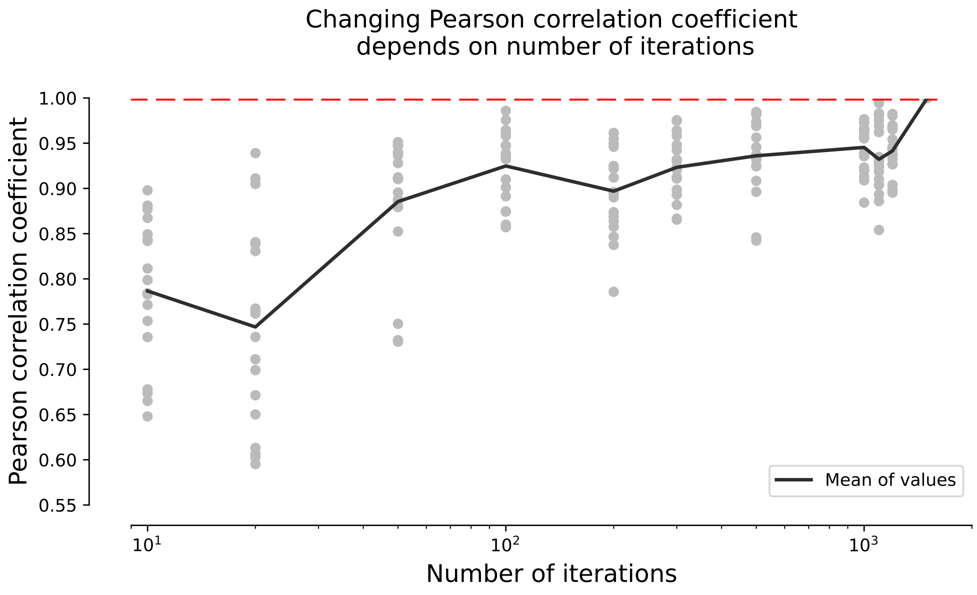


Supplementary Figure 4: Convergence of the relative model score with an increase in the number of shuffle iterations
